# Fundamental frequency predominantly drives talker differences in auditory brainstem responses to continuous speech

**DOI:** 10.1101/2024.07.12.603125

**Authors:** Melissa J. Polonenko, Ross K. Maddox

## Abstract

Deriving human neural responses to natural speech is now possible, but the responses to male- and female-uttered speech have been shown to differ. These talker differences may complicate interpretations or restrict experimental designs geared toward more realistic communication scenarios. This study found that when a male and female talker had the same fundamental frequency, auditory brainstem responses (ABRs) were very similar. Those responses became smaller and later with increasing fundamental frequency, as did click ABRs with increasing stimulus rates. Modeled responses suggested that the speech and click ABR differences were reasonably predicted by peripheral and brainstem processing of stimulus acoustics.

## 1. Introduction

Aural communication in daily life involves listening to and parsing out the continuously dynamic spectral-temporal content that comprises natural speech. Only recently have researchers been able to use such naturally dynamic speech to investigate auditory neural processing through responses derived from electroencephalography (EEG) or magnetoencephalography. With several of these different new methods, however, the responses to female talkers seem to be smaller than to male talkers (Canneyt et al., 2021; Commuri et al., 2023; Easwar et al., 2021, 2022; Maddox and Lee, 2018; Polonenko and Maddox, 2021; Saiz-Alía et al., 2019; Saiz-Alia and Reichenbach, 2020), which can have implications/consequences for experiments requiring multiple talkers to more closely resemble real-life scenarios.

It is well known that higher stimulation rates decrease amplitude and increase latency for auditory responses to click and toneburst stimuli (e.g., Burkard et al., 1990; Burkard and Hecox, 1983; Don et al., 1977; Jiang et al., 2009; Polonenko and Maddox, 2022). For naturally uttered speech, male talkers tend to have lower fundamental frequencies (f0, Hillenbrand et al., 1995), a result of typically more massive folds vocal folds coming together and reopening at slower rates to produce voiced speech. This essentially gives a lower rate of “glottal pulses” corresponding to the lower pitch. Therefore, it’s reasonable to expect that differing f0, rather than gender itself, may be driving the talker differences in auditory responses. Recent studies do suggest that f0 contributes to the male/female differences in envelope-following responses, brainstem responses, and cortical temporal response functions (Canneyt et al., 2021; Easwar et al., 2021, 2022; Saiz-Alia and Reichenbach, 2020), but this effect was studied using talkers that also differ in characteristics besides f0.

In this study, the f0 of two audiobook stimuli—one narrated by a female, the other by a male—is manipulated in order to systematically test the effect of increasing f0 on auditory brainstem responses (ABRs) to continuous speech. The f0 effect on speech ABRs is also compared to the stimulation rate effect on click ABRs with clicks set to the same rate as the speech f0s. To study these f0/rate effects, the “peaky speech” method is used because the speech ABRs show canonical morphologies that reflect activity from distinct neurogenerators from the auditory nerve to rostral brainstem and can be derived in a similar method to click ABRs (Polonenko and Maddox, 2021). As shown in Figure 1a, the phase structure of the continuous natural speech is re-synthesized to make the speech waveform as click-like as possible (Figure 1a, left column) while preserving the spectral-temporal properties (Figure 1a, right column spectrograms). This is done by re-constructing the speech phase structure from the glottal pulse trains derived from the f0, which themselves act like the pulses in click trains. Previous work with continuous peaky speech from narrated audiobooks also shows the male/female talker difference for speech ABRs (Figure 1b, left column)(Polonenko and Maddox, 2021).

This study aims to: 1) replicate the male/female talker effect with each at their natural f0, 2) systematically determine if f0 is the main driver of this talker difference, and 3) evaluate if the f0 effect resembles the click rate effect. ABRs to the same stimuli are also modeled to explore whether any differences in ABRs to speech and clicks are primarily acoustically driven and due to peripheral and brainstem auditory processing (as the model does not include any cortical contributions). Computational modeling has been previously used to simulate ABRs and explore stimulus differences (e.g., Dau, 2003; Stoll and Maddox, 2023; Temboury-Gutierrez et al., 2024).

**Figure 1.**
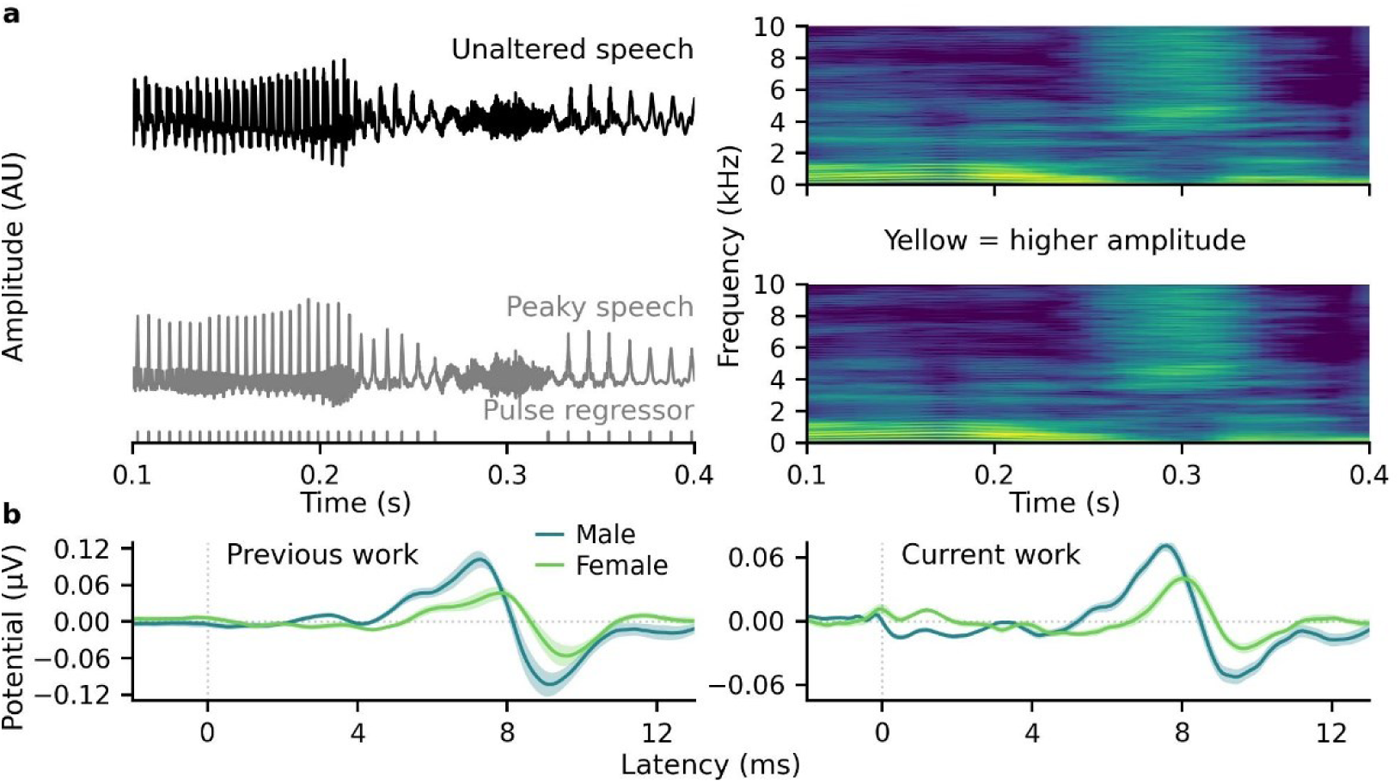
Peaky speech stimuli and evoked ABRs. (a) The peaky speech waveform (bottom left) is more “click-like” than the unaltered waveform (top left), but their spectrograms are essentially the same (right) because only the phase was altered. (b) Average ABRs (areas show ± SE) measured in previous work (left, n = 11) and the current study (right, n = 15) are larger and earlier for male-than female-narrated speech. Data for previous work adapted from (Polonenko and Maddox, 2021).

## 2. Methods

### 2.1 Participants

Data were collected under a protocol approved by the University of Rochester Research Subjects Review Board (#1227). All participants gave informed consent and were compensated for their time. The 15 participants were aged 19–35 years with a mean ± SD age of 24.1 ± 6.1 years, and included 5 males and 10 females. Audiometric screening confirmed participants had normal hearing in both ears, defined as thresholds ≤20 dB HL from 250–8000 Hz. All participants identified English as their primary language.

### 2.2 Stimuli

An hour of each of the same audiobooks was used as before (Polonenko and Maddox, 2021): *The Alchemyst* read by a male narrator with an average fundamental frequency (f0) of 123 Hz (Scott, 2007) and *A Wrinkle in Time* read by a female narrator with an f0 of 183 Hz (L’Engle, 2012). The pitch of each narrator’s audio was shifted using PRAAT based on a factor determined by the semitone differences in pitch between the narrators (Boersma and Weenink, 2018): unshifted, the f0 of the other narrator, and the geometric mean of the two narrators (i.e., 150 Hz), for a total of 6 conditions. Audio processing included resampling to 48 kHz, truncating silences to ≤0.5 s, splicing into 10 s segments with 0.03 s cosine fade-in/out, shifting the mean f0, then re-synthesizing into broadband peaky speech as done in prior work (Polonenko and Maddox, 2021). The re-synthesized voiced portions of speech were combined with the unvoiced portions of the original speech. Polarity alternated between speech segments within each trial to limit stimulus artifacts in the EEG.

Click stimuli were also created as before (Bachmann et al., 2024; Maddox and Lee, 2018) but with stimulus rates corresponding to the shifted speech f0s (123, 150, 183 stim / s). A total of 150 unique 1 s epochs were created with a pseudorandom Poisson process controlling the timing of the click trains. An inverted version of each epoch was presented in sequence to counter-phase the stimuli to help mitigate stimulus artifact (i.e., Epoch A^+^A^-^B^+^B^-^, where the letter indicates the timing sequence and the +/-indicates the phase). These 300 epochs were concatenated into 10 s segments to utilize the same soundcard buffer for both types of stimuli.

The inter-stimulus intervals (ISIs) for each frequency shift (123, 150, 183 Hz) and stimulus (clicks, male narrator, female narrator) are compared in Figure 2. For speech, the ISI is considered the time between adjacent glottal pulses. The click ISIs follow the Poisson distribution used to create the randomized click trains, whereas the speech ISIs show most ISIs close to the inverse of the mean f0.

**Figure 2.**
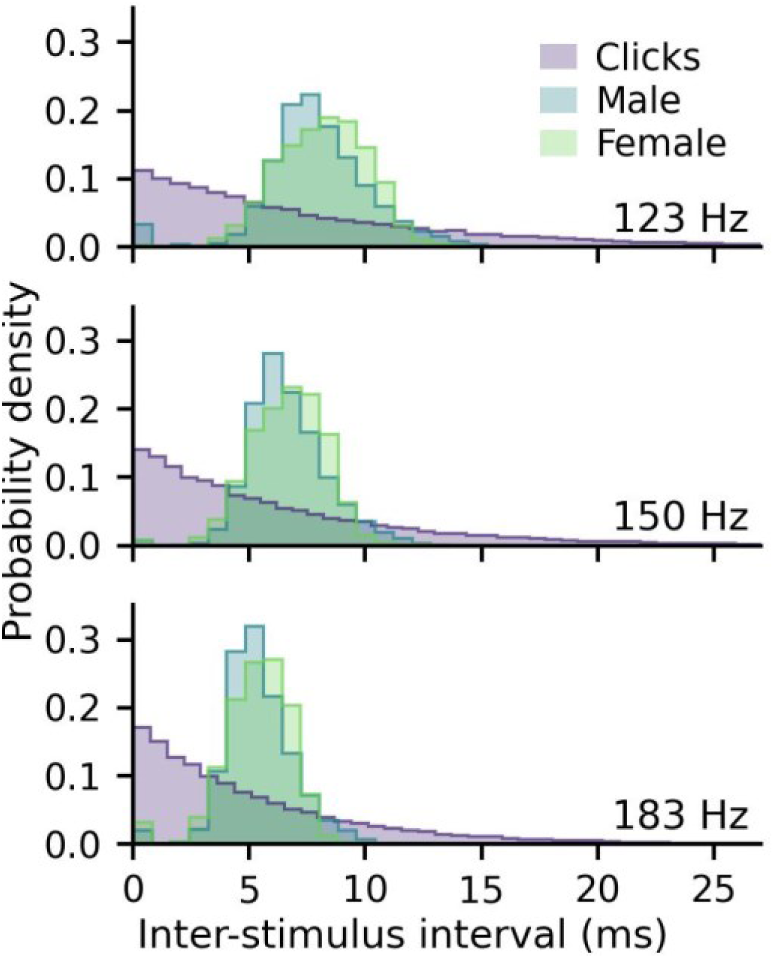
Stimulus ISIs. The densities of inter-stimulus intervals (ISIs) for speech and clicks were set to average f0s/rates of 123, 150, and 183 Hz.

### 2.3 Data collection

Participants listened to 15 minutes of clicks (5 minutes per rate) and then 2 hours of speech (20 minutes per narrator-f0 condition) while reclining and resting or watching silent subtitled videos in a darkened sound booth. Stimulus presentation was randomized and controlled by a custom python script built on the expyfun module (Larson et al., 2014). Stimuli were played diotically at 65 dB (pe)SPL over Etymotic Research ER-2 insert earphones hung from the ceiling and plugged into an RME Babyface Pro soundcard at 48 kHz.

EEG potentials were recorded with a sampling rate of 10 kHz using BrainVision’s PyCorder software with two EP-Preamp preamplifiers connected to the actiCHamp amplifier. Two bipolar channels were recorded with Fz referenced to each earlobe and the ground placed at Fpz. Triggers for synchronizing the audio and EEG were created by the soundcard’s optical digital output and converted to TTL voltages using a custom trigger box (Maddox, 2020) before being sent to the EEG amplifier. These triggers precisely denoted the start and end of each trial to correct drift between the soundcard and amplifier clocks. The 0.9 ms earphone tubing delay was corrected during preprocessing.

### 2.4 Preprocessing and ABR derivation

Preprocessing and analysis were done with custom python scripts that used the mne module (Gramfort et al., 2013). Raw EEG data were filtered between 150 and 2000 Hz using a first-order causal Butterworth bandpass filter to remove slow drift in the signal and optimize ABR morphology, and with 5 Hz wide second-order infinite impulse response notch filters at odd multiples of 60 Hz to remove electrical line noise.

Epochs were created for each 10 s trial, including 1 s before and after the stimulus for 12 s of EEG per trial. Similarly, the impulse sequence for the trial clicks or speech glottal pulses was used to create a regressor pulse train that was zero-padded with 1 s before and after to give a 12s pulse train. The EEG and pulse trains were then used to derive ABRs as described previously for clicks using cross-correlation (Maddox and Lee, 2018) and for peaky speech using deconvolution (Polonenko and Maddox, 2021). For efficiency, cross-correlations and deconvolutions were computed in the frequency domain. A Bayesian-like weighted average was used to improve signal-to-noise ratio (SNR), as in our previous work (Elberling and Wahlgreen, 1985; Polonenko and Maddox, 2019, 2021, 2022). Each trial segment of EEG was weighted by its inverse variance divided by the sum of the inverse variances of all trials for that condition. ABRs derived from the two channels were averaged together to further increase SNR.

### 2.5 Modeled responses

Simulated ABRs were derived for each stimulus using well-known computational models of the auditory nerve and periphery that account for acoustics and some of the non-linearities of the auditory system (Rudnicki et al., 2015; Verhulst et al., 2018; Zilany et al., 2014). The EEG was simulated for waves I, III and V using the framework described by Verhulst et al. (2018) but with the peripheral model by Zilany et al. (2014) due to computation constraints. The default scale and latency shift for each wave were adjusted with a grid search until the modeled ABR for the lowest rate clicks matched that of the grand average measured ABR. The simulated EEG of each model was then cross-correlated (clicks) or deconvolved (speech) with the respective pulse trains to derive the modeled ABRs.

### 2.6 Statistical Analyses

ABR waveform morphologies from 2 to 12 ms were compared using the Pearson correlation between male- and female-narrated speech at each f0 and for each participant. These correlation coefficients were compared to the null set of correlations for the ABRs split into even and odd trials with equal number of trials of male- and female-narrated speech using the Wilcoxon signed-rank test, as done before (Polonenko and Maddox, 2021). False discovery rate (FDR, Benjamini and Hochberg, 1995) was used to correct p-values for family-wise errors.

The peak was picked for the most prominent wave V and its following trough to calculate the peak-to-trough amplitude. These chosen peaks are shown on ABRs for each participant in Supplementary Material Figures 1–3 for the three frequency shifts respectively. The ratios of wave V for male/female speech across participants were compared to those of the previous study using a Mann-Whitney U test. Changes in wave V peak amplitudes and latencies were compared using linear mixed-effects regression with a random intercept per participant and fixed effects of stimulus, rate/f0, and their two-way interaction. Mixed models were performed in RStudio with the lme4 and lmerTest packages (Bates et al., 2014; Kuznetsova et al., 2017; RStudio Team, 2021; Russell V. Lenth, 2022).

## 3. Results

### 3.1 The talker effect on speech ABRs is replicated at natural f0

The grand average ABRs to the natural (unshifted) male- and female-narrated peaky speech are shown in Figure 1b. They showed similar canonical morphologies, but the responses to male-narrated speech were earlier and 1.80 ± 0.14 (mean ± SE) times larger than for female-narrated speech. The male/female wave V difference was similar to the 1.98 ± 0.10 ratio from previous ABRs using the same narrated stories (Polonenko and Maddox, 2021) (Mann-Whitney U test, M = 118, p = 0.069).

**Figure 3.**
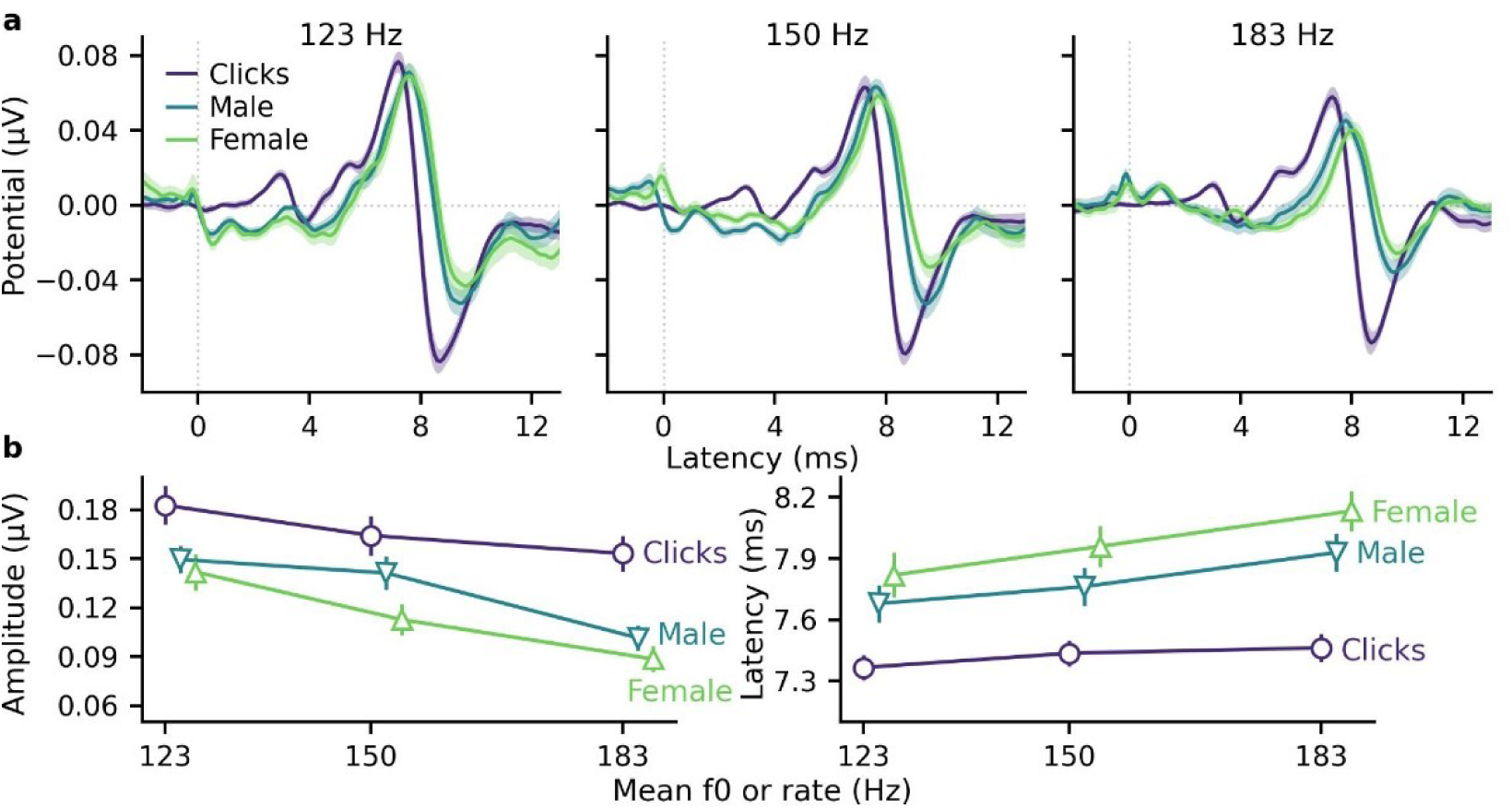
Comparison of measured ABRs evoked by stimuli with similar average rate/f0s. (a) Average ABRs (areas show ± SE) to male- and female-narrated speech with the same mean f0 were similar to each other but different to clicks with the same mean rate. (b) Mean ± SE wave V peak amplitudes (left) decreased, and latencies (right) increased with increasing rate/f0 and were the smallest and latest for the female-narrated speech. Details of the mixed effects models are provided in Supplementary Material Tables 1 and 2.

### 3.2 Measured ABRs to the same mean f0 or rate are similar, independent of talker

Figure 3a demonstrates that shifting the narrators’ speech to the same mean f0 gave similar ABRs, although responses were still slightly larger and earlier for the male speech. Over latencies 2–12 ms, the median correlation coefficients between the male/female responses for 123, 150, and 183 Hz were high at 0.89, 0.89, and 0.80 respectively, which were only significantly lower than the median null split-half correlations for the lowest f0 (123, 150, 183 Hz: medians 0.96, 0.93, 0.83; Wilcoxon signed-rank test W=10, 28, 44; FDR-adjusted p: 0.008, 0.109, 0.389 respectively). As shown in Figure 3b and the mixed models in Supplemental Tables 1–2, male speech wave V peak amplitudes were larger (16.3 ± 5.1 nV, p = 0.002) and earlier (-0.17 ± 0.03 ms, p < 0.001) than the female peaks, especially at 150 Hz f0, but were more similarly sized for 123 and 183 Hz f0s. Peaks became smaller and later with increasing f0 (both p < 0.001), but the overall rates of change were similar for both narrators (slope differences: 0.07 ± 0.21 nV/Hz and 0.001 ± 0.001 ms/Hz, p > 0.419). Thus, matching the mean f0 reduced much of the previous differences between male/female ABRs (Figure 1b), but some small differences remained.

Previous work has also shown that continuous speech ABRs are smaller and later than click ABRs, but click ABRs are mostly done with rates much lower than the f0s of speech, which adds a confound. Figure 3a shows that using a mean click rate similar to the speech f0s gives more proportionally/similarly sized ABRs, but the click ABRs are still earlier, larger and have more distinct morphology (i.e., waves I and III) than the speech ABRs. The click wave V peak amplitudes and latencies shown in Figure 3b were larger (35.1 ± 5.1 nV, p < 0.001) and earlier (-0.35 ± 0.03 ms, p < 0.001) than to speech, but the changes with increasing click rate were similar to those with f0 for the speech ABRs (slope differences: 0.32 ± 0.21 nV/Hz, 0.003 ± 0.001 ms/Hz; both p < 0.001).

### 3.3 Modeled ABRs show similar trends to measured ABRs for amplitude but not latency

**Figure 4.**
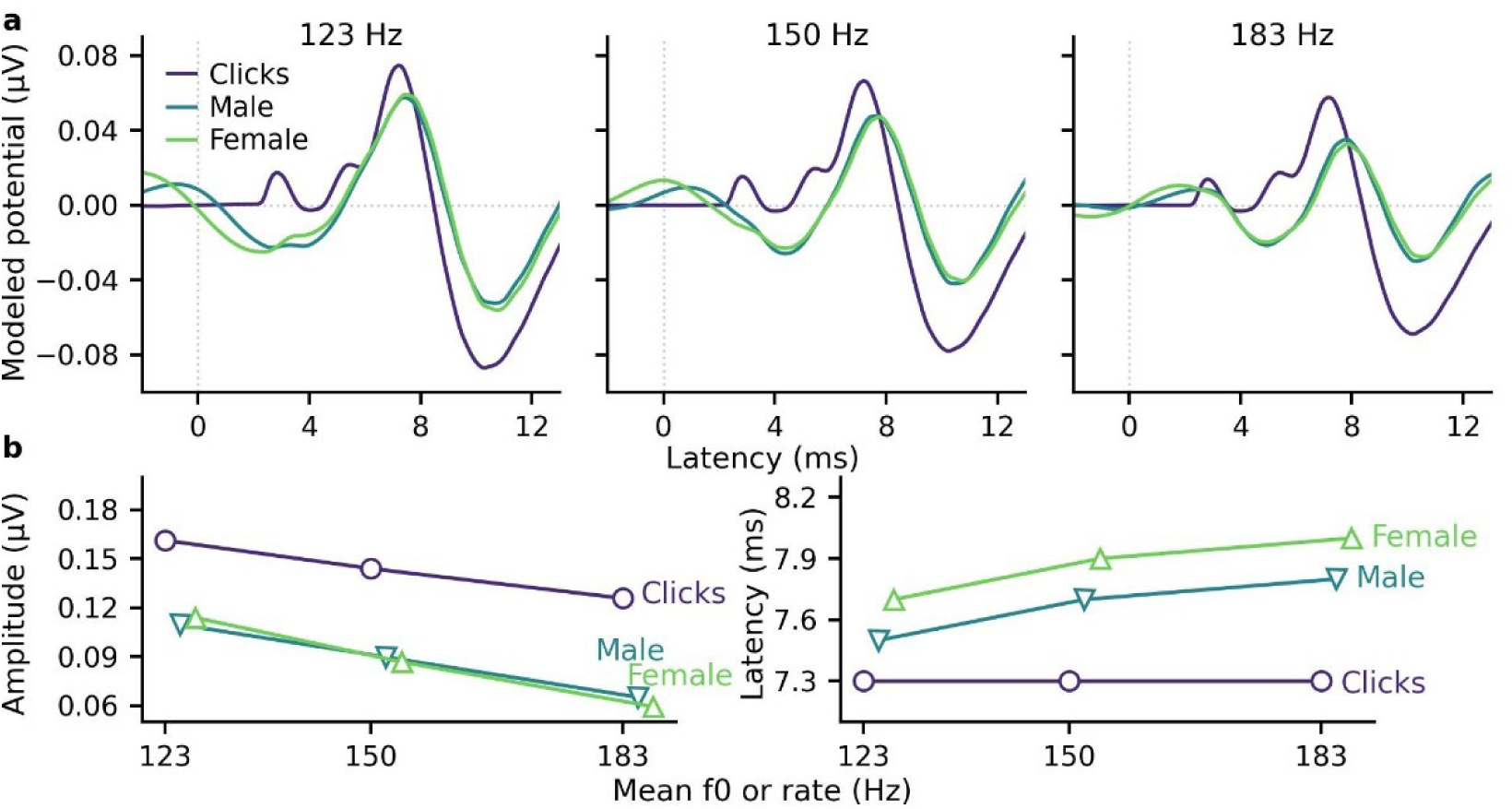
Comparison of modeled ABRs to stimuli with similar average rate/f0s. (a) Modeled ABRs to male- and female-narrated speech with the same mean f0 were very similar but had poorer morphology and smaller, later wave V peaks than to clicks with the same mean rate. (b) Wave V peak amplitudes decreased (left) and latencies (right) increased with increasing rate/f0 except the pattern did not hold for click wave V latencies.

Next, ABRs were modeled for the three stimuli to determine whether acoustics and peripheral and brainstem processing accounted for the stimulus and f0/rate trends observed in the measured ABRs. The modeled ABR waveforms in Figure 4a, and the wave V peak amplitudes and latencies in Figure 4b, replicated the main changes for speech ABRs, except the small male/female amplitude differences were not replicated. Modeled click responses were larger and earlier, but they did not show any latency changes with increasing click rate the measured ABRs showed.

## 4. Discussion

This study systematically evaluated the effect of f0 on speech ABRs to a male and female talker and compared these effects to rate effects on click ABRs. Speech ABRs decreased in amplitude and increased in latency with increasing f0 with similar slopes for both narrators. Once set to the same mean f0, ABRs to an originally male and female talker were essentially similar, with only small differences remaining. With a similar mean click rate, the click ABRs were still larger and earlier than the speech ABRs but to a lesser extent than previously observed. Modeling replicated the changes for speech and the differences between speech and clicks, suggesting that the differences are largely due to acoustical peripheral and brainstem processing and not higher-order categorization.

### 4.1 The f0 accounted for much of the talker effects on ABRs

This study replicated the male/female talker effects on continuous speech ABRs (Figure 1b) but also demonstrated that f0 accounted for a large portion of these ABR differences (Figure 3a). While important to confirm, this result is not surprising, given the well-known rate effects for ABRs to brief stimuli (e.g., Burkard et al., 1990; Burkard and Hecox, 1983; Don et al., 1977; Jiang et al., 2009; Polonenko and Maddox, 2022), and the lower amplitudes for talkers with higher f0s in other auditory responses using different methods with continuous speech (Canneyt et al., 2021; Easwar et al., 2021, 2022; Maddox and Lee, 2018; Saiz-Alía et al., 2019; Saiz-Alia and Reichenbach, 2020). What this study adds is the systematic confirmation of f0 driving much of the ABR differences by keeping the same talkers while only adjusting the f0. Furthermore, the speech ABRs showed changes with increasing f0 similar to click ABRs with increasing stimulation rate. This suggests neural adaptation to the glottal pulse rate (i.e., f0) akin to stimulation rate, which is consistent with f0 rather than a categorical gender effect driving the narrator-dependent ABR differences.

### 4.2 Talker differences remain after matching mean f0

Even when f0 was equal, the talker with the originally lower f0 had slightly larger and earlier responses, particularly for the mid f0 rate (Figure 3). These remaining response differences could result from differences between the two talkers, but no claim can be made about gender with only one sample of each. A variety of factors could explain the differences, and talker identity, gender, or other talker characteristics may have small effects on the ABRs. The differences are unlikely due to different text because the ABRs split into even/odd trials also had different text but high correlations (>0.9, see section 3.2). However, the rate of f0 change within the narrated text has been shown to affect neural tracking in cortical responses in addition to f0 (Canneyt et al., 2021), and narrators used in this study could have mimicked different character voices in a way that differentially impacted the rate of local f0 changes during the story. Even though mean f0 was matched the ISI distributions slightly differed (Figure 2), but the rate of f0 change was not evaluated herein. Pilot studies in the authors’ labs also suggest that ABRs can vary between talkers of the same gender and similar f0 statistics in ways that are hard to predict.

### 4.3 Speech ABRs are more similar to click ABRs when the mean f0 matches the mean rate

Previous work noted that speech ABRs are smaller than reported values for click ABRs (Maddox and Lee, 2018; Polonenko and Maddox, 2021), which is at least partially due to the dynamically changing acoustics of continuous speech. Click ABRs, however, are usually evoked with lower stimulation rates than the typical speech f0s (>60 Hz), confounding the comparison. The lower rates are necessary for periodic stimulation due to the response window, but are also generally done for randomized stimuli (Bachmann et al., 2024; e.g., Maddox and Lee, 2018; Polonenko and Maddox, 2019, 2022), except for specific studies with high-rate clicks (e.g., Valderrama et al., 2012). In this study, matching the mean click rate to the mean speech f0s brought the click and speech ABRs into a similar amplitude range, suggesting that stimulation rate may partially account for the previously described click-speech ABR differences.

Differences remained despite matching mean rate/f0 of the clicks and speech: the speech ABRs were still smaller and had less distinct earlier component waves than the click ABRs (Figure 3). These stimulus-related differences could be due to a variety of factors. Although the peaky speech method evokes more synchronous activity with the altered phase structure of the speech (Polonenko and Maddox, 2021), there are still dynamic variations in the speech that are not present in the click trains. Clicks are very transient and evoke highly synchronous responses, giving distinct component waves even at higher rates matching speech f0s. Clicks are also broadband, with a flat frequency response up to 10 kHz, containing more high-frequency energy than the sloping amplitude spectrum of natural speech stimuli.

Another possible reason for the differences may be the underlying ISI distributions used to create the clicks. In this study, a pseudorandom Poisson process was used to create randomized click trains as in prior work (Bachmann et al., 2024; Maddox and Lee, 2018). Although the mean rates matched mean f0s, the exponential ISI distribution characteristic of a Poisson process (Figure 2) differed from the speech ISI distributions and may account for some of the ABR differences between the two types of stimuli. Future work could use the same glottal pulse trains as the speech to create the click stimuli to evaluate whether click-speech differences remain when the ISI distributions also match. Regardless, matching mean rate/f0 seems to play a partial role in reducing some of the previously observed size differences for click and speech ABRs.

### 4.4 Modeled ABRs reasonably approximate stimulus-f0 effects, but not click latencies or small talker differences

Modeled ABRs roughly approximated the main speech and click ABR differences (Figure 4), with a few exceptions. Both the f0 effect on speech ABRs and the speech/click ABR differences were replicated, further providing evidence that subcortical processing of stimulus acoustics over a categorical distinction – which is not considered by the computational model – primarily underlie the natural (unshifted) talker differences and speech/click stimulus differences in the ABRs.

However, the model is not fully accurate and did not predict the remaining talker differences after matching f0 or the latency changes in click ABRs with increasing rate. These limitations of the computational model are consistent with other studies that also found appropriate predictions of amplitude but not latency changes with stimulus level changes in click ABRs (e.g., Dau, 2003; Temboury-Gutierrez et al., 2024). They suggested that the limitations could be due to an oversimplification of neural processing in the brainstem model, which assumes constant delays through the brainstem that may not be accurate for broadband or complex stimuli. Cortical processes that are not captured by the computation model may also contribute to the remaining talker differences. Nevertheless, computational modeling provides a reasonable prediction of the main stimulus parametric effects on both speech and click ABR amplitudes that can be useful when planning human experiments with different stimuli and for evaluating the contributions of acoustics versus categorizations.

## 5. Conclusion

Speech ABRs show decreased amplitude and increased latency with increasing f0. The same is true of clicks for increasing stimulus rate. Once stimuli are set to the same f0/rate, ABRs to speech and clicks are more similar than previously reported, although click ABRs remain larger and earlier than the speech ABRs, and small differences remain between ABRs for the two narrators. Modeled responses mostly replicated the main effects for both clicks and speech. These results suggest that the previous stimulus effects, particularly between male and female narrators, are largely driven by differences in f0 (or stimulus rate, for clicks), with the subtle remaining narrator differences likely due to several variables not systematically evaluated here.

## Acknowledgments

This work was supported by NIDCD R01DC017962. The authors thank Yathida Melody Anankul for assistance with recruitment and data collection.

## Author Declarations

### Conflict of Interest

The authors have no conflicts to disclose.

### Ethics Approval

Written informed consent was obtained from all participants before the experiment according to protocol #1227 approved by the University of Rochester Research Subjects Review Board.

## Data Availability

The data that support the findings of this study are available within the article and its Supplementary Material, as well as openly available in OpenNeuro, reference number ds005340.

## Notes

### Competing Interest Statement

The authors have declared no competing interest.

## References and Links

Bachmann, F. L., Kulasingham, J. P., Eskelund, K., Enqvist, M., Alickovic, E., and Innes-Brown, H. (2024). “Extending Subcortical EEG Responses to Continuous Speech to the Sound-Field,” Trends in Hearing, 28, 23312165241246596. doi:10.1177/23312165241246596

Bates, D., Maechler, M., Bolker, B. M., and Walker, S. (2014). lme4: Linear mixed-effects models using Eigen and S4. ArXiv e-print; submitted to Journal of Statistical Software,.

Benjamini, Y., and Hochberg, Y. (1995). “Controlling the False Discovery Rate: A Practical and Powerful Approach to Multiple Testing,” Journal of the Royal Statistical Society. Series B (Methodological), 57, 289–300.

Boersma, P., and Weenink, D. (2018). “Praat: doing phonetics by computer.,” Retrieved from http://www.praat.org/

Burkard, R., and Hecox, K. (1983). “The effect of broadband noise on the human brainstem auditory evoked response. I. Rate and intensity effects,” The Journal of the Acoustical Society of America, 74, 1204–1213. doi:10.1121/1.390024

Burkard, R., Shi, Y., and Hecox, K. E. (1990). “A comparison of maximum length and Legendre sequences for the derivation of brain-stem auditory-evoked responses at rapid rates of stimulation,” The Journal of the Acoustical Society of America, 87, 1656–1664. doi:10.1121/1.399413

Canneyt, J. V., Wouters, J., and Francart, T. (2021). “Neural tracking of the fundamental frequency of the voice: The effect of voice characteristics,” European Journal of Neuroscience, 53, 3640–3653. doi:10.1111/ejn.15229

Commuri, V., Kulasingham, J. P., and Simon, J. Z. (2023). “Cortical responses time-locked to continuous speech in the high-gamma band depend on selective attention,” Front. Neurosci., doi: 10.3389/fnins.2023.1264453. doi:10.3389/fnins.2023.1264453

Dau, T. (2003). “The importance of cochlear processing for the formation of auditory brainstem and frequency following responses,” The Journal of the Acoustical Society of America, 113, 936–950. doi:10.1121/1.1534833

Don, M., Allen, A. R., and Starr, A. (1977). “Effect of Click Rate on the Latency of Auditory Brain Stem Responses in Humans,” Ann Otol Rhinol Laryngol, 86, 186–195. doi:10.1177/000348947708600209

Easwar, V., Boothalingam, S., and Regan, F. (2021). “Fundamental frequency-dependent changes in vowel-evoked envelope following responses,” Hearing Research, doi: 10.1016/j.heares.2021.108297. doi:10.1016/j.heares.2021.108297

Easwar, V., Purcell, D., Eeckhoutte, M. V., and Aiken, S. J. (2022). “The Influence of Male- and Female-Spoken Vowel Acoustics on Envelope-Following Responses,” Semin Hear, 43, 223–239. doi:10.1055/s-0042-1756165

Elberling, C., and Wahlgreen, O. (1985). “Estimation of auditory brainstem response, ABR, by means of Bayesian inference,” Scand Audiol, 14, 89–96. doi:10.3109/01050398509045928

Gramfort, A., Luessi, M., Larson, E., Engemann, D. A., Strohmeier, D., Brodbeck, C., Goj, R., et al. (2013). “MEG and EEG data analysis with MNE-Python,” Front. Neurosci., doi: 10.3389/fnins.2013.00267. doi:10.3389/fnins.2013.00267

Hillenbrand, J., Getty, L. A., Clark, M. J., and Wheeler, K. (1995). “Acoustic characteristics of American English vowels,” J Acoust Soc Am, 97, 3099– 3111. doi:10.1121/1.411872

Jiang, Z. D., Wu, Y. Y., and Wilkinson, A. R. (2009). “Age-related changes in BAER at different click rates from neonates to adults,” Acta Paediatr., 98, 1284– 1287. doi:10.1111/j.1651-2227.2009.01312.x

Kuznetsova, A., Brockhoff, P. B., and Christensen, R. H. B. (2017). “lmerTest Package: Tests in Linear Mixed Effects Models,” Journal of Statistical Software, 82, 1–26. doi:10.18637/jss.v082.i13

Larson, E., McCloy, D., Maddox, R., and Pospisil, D. (2014). “expyfun: Python experimental paradigm functions, version 2.0.0.,” doi:10.5281/zenodo.11640

L’Engle, M. (2012). *A Wrinkle in Time. Audiobook*, Listening Library, Available: https://www.booksontape.com/book/96850/a-wrinkle-in-time/, (date last viewed: 26-Jan-21). Retrieved January 26, 2021, from https://www.booksontape.com/book/96850/a-wrinkle-in-time/

Maddox, R. K. (2020). “S/Plitter: Hardware and Firmware for Converting Digital Audio to TTL Triggers.,” Retrieved from 10.5281/zenodo.10802516

Maddox, R. K., and Lee, A. K. C. (2018). “Auditory Brainstem Responses to Continuous Natural Speech in Human Listeners,” eNeuro, 5, ENEURO.0441-17.2018. doi:10.1523/ENEURO.0441-17.2018

Polonenko, M. J., and Maddox, R. K. (2019). “The Parallel Auditory Brainstem Response,” Trends Hear, 23, 2331216519871395. doi:10.1177/2331216519871395

Polonenko, M. J., and Maddox, R. K. (2021). “Exposing distinct subcortical components of the auditory brainstem response evoked by continuous naturalistic speech,” (T. Reichenbach, A. J. King, T. Reichenbach, and J. Z. Simon, Eds.) eLife, 10, e62329. doi:10.7554/eLife.62329

Polonenko, M. J., and Maddox, R. K. (2022). “Optimizing Parameters for Using the Parallel Auditory Brainstem Response to Quickly Estimate Hearing Thresholds,” Ear Hear, 43, 646–658. doi:10.1097/AUD.0000000000001128

RStudio Team (2021). “RStudio: Integrated Development Environment for R.,” Retrieved from http://www.rstudio.com/

Rudnicki, M., Schoppe, O., Isik, M., Völk, F., and Hemmert, W. (2015). “Modeling auditory coding: from sound to spikes,” Cell Tissue Res, 361, 159–175. doi:10.1007/s00441-015-2202-z

Russell V. Lenth (2022). “emmeans: Estimated Marginal Means, aka Least-Squres Means.,” Retrieved from https://CRAN.R-project.org/package=emmeans

Saiz-Alía, M., Forte, A. E., and Reichenbach, T. (2019). “Individual differences in the attentional modulation of the human auditory brainstem response to speech inform on speech-in-noise deficits,” Scientific Reports, 9, 1–10. doi:10.1038/s41598-019-50773-1

Saiz-Alia, M., and Reichenbach, T. (2020). “Computational modeling of the auditory brainstem response to continuous speech,” J. Neural Eng., doi: 10.1088/1741-2552/ab970d. doi:10.1088/1741-2552/ab970d

Scott, M. (2007). *The Alchemyst: the secrets of the immortal Nicholas Flamel, Book 1*. *Audiobook*, Listening Library, Available: https://www.booksontape.com/book/163186/the-alchemyst/. Retrieved from https://www.booksontape.com/book/163186/the-alchemyst/

Stoll, T. J., and Maddox, R. K. (2023). “Enhanced Place Specificity of the Parallel Auditory Brainstem Response: A Modeling Study,” Trends Hear, 27, 23312165231205719. doi:10.1177/23312165231205719

Temboury-Gutierrez, M., Encina-Llamas, G., and Dau, T. (2024). “Predicting early auditory evoked potentials using a computational model of auditory-nerve processing,” The Journal of the Acoustical Society of America, 155, 1799– 1812. doi:10.1121/10.0025136

Valderrama, J. T., Alvarez, I., de la Torre, A., Carlos Segura, J., Sainz, M., and Luis Vargas, J. (2012). “Recording of auditory brainstem response at high stimulation rates using randomized stimulation and averaging,” The Journal of the Acoustical Society of America, 132, 3856–3865. doi:10.1121/1.4764511

Verhulst, S., Altoè, A., and Vasilkov, V. (2018). “Computational modeling of the human auditory periphery: Auditory-nerve responses, evoked potentials and hearing loss,” Hearing Research, Computational models of the auditory system, 360, 55–75. doi:10.1016/j.heares.2017.12.018

Zilany, M. S. A., Bruce, I. C., and Carney, L. H. (2014). “Updated parameters and expanded simulation options for a model of the auditory periphery,” The Journal of the Acoustical Society of America, 135, 283–286. doi:10.1121/1.4837815

