## Supplementary Material for "Fundamental frequency predominantly drives talker differences in auditory brainstem responses to continuous speech"

**Supplemental Materials**

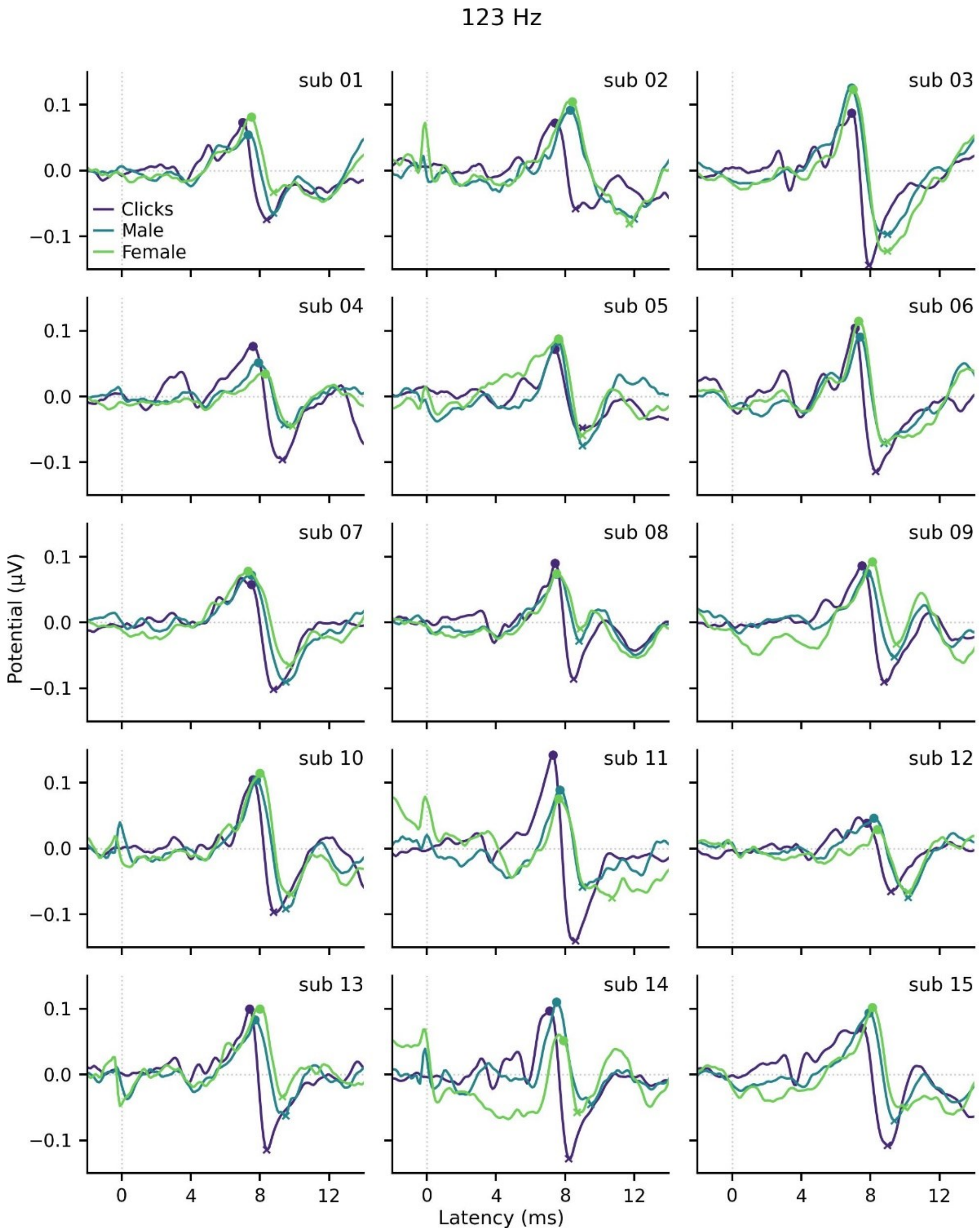


Supplemental Figure 1. Individual participant ABRs are shown for clicks, male- and female-narrated speech with a mean rate or f0 of 123 Hz. Wave V peak latency (circle) and trough (x) are marked, which were used to calculate amplitude.

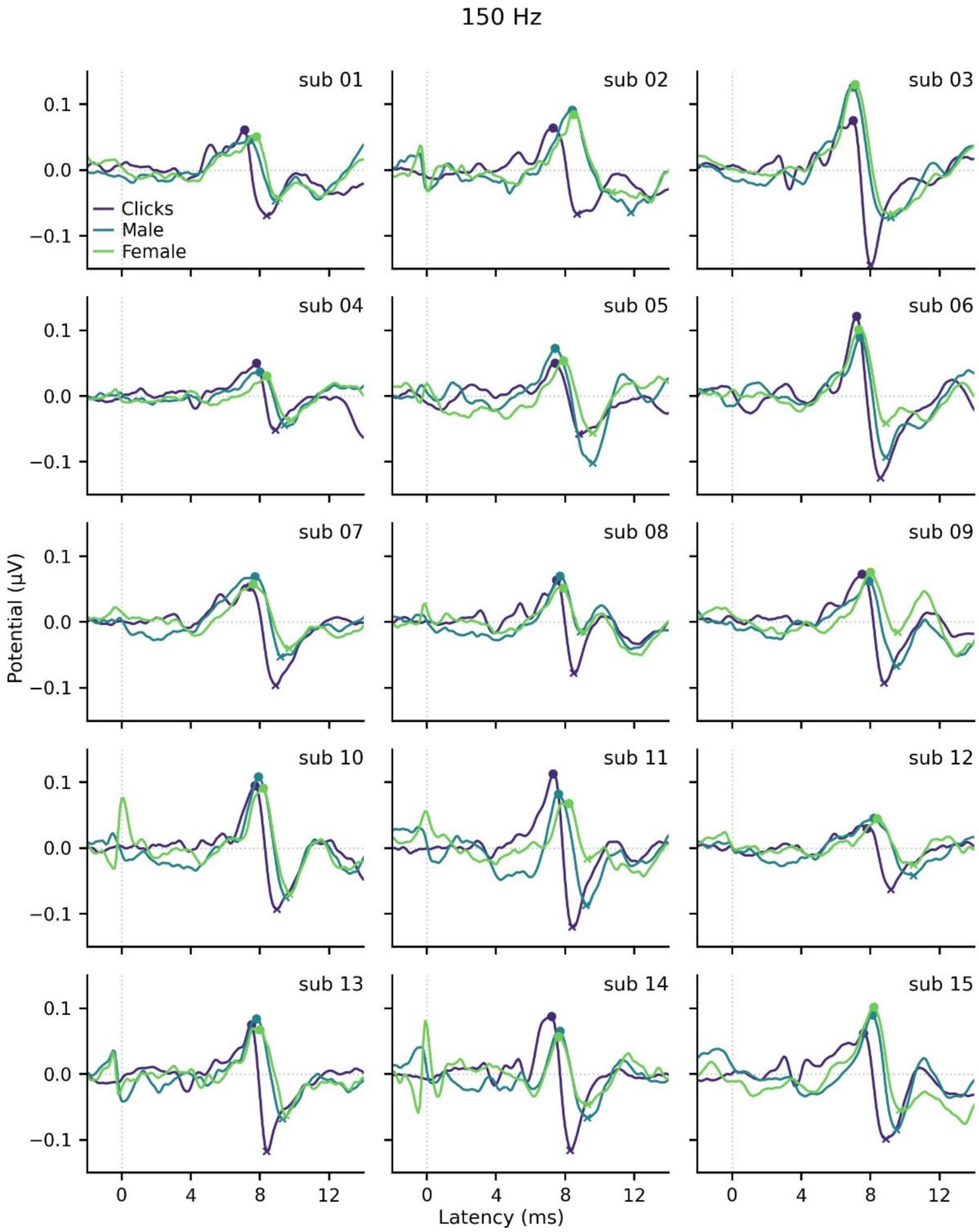


Supplemental Figure 2. Individual participant ABRs are shown for clicks, male- and female-narrated speech with a mean rate or f0 of 150 Hz. Wave V peak latency (circle) and trough (x) are marked, which were used to calculate amplitude.

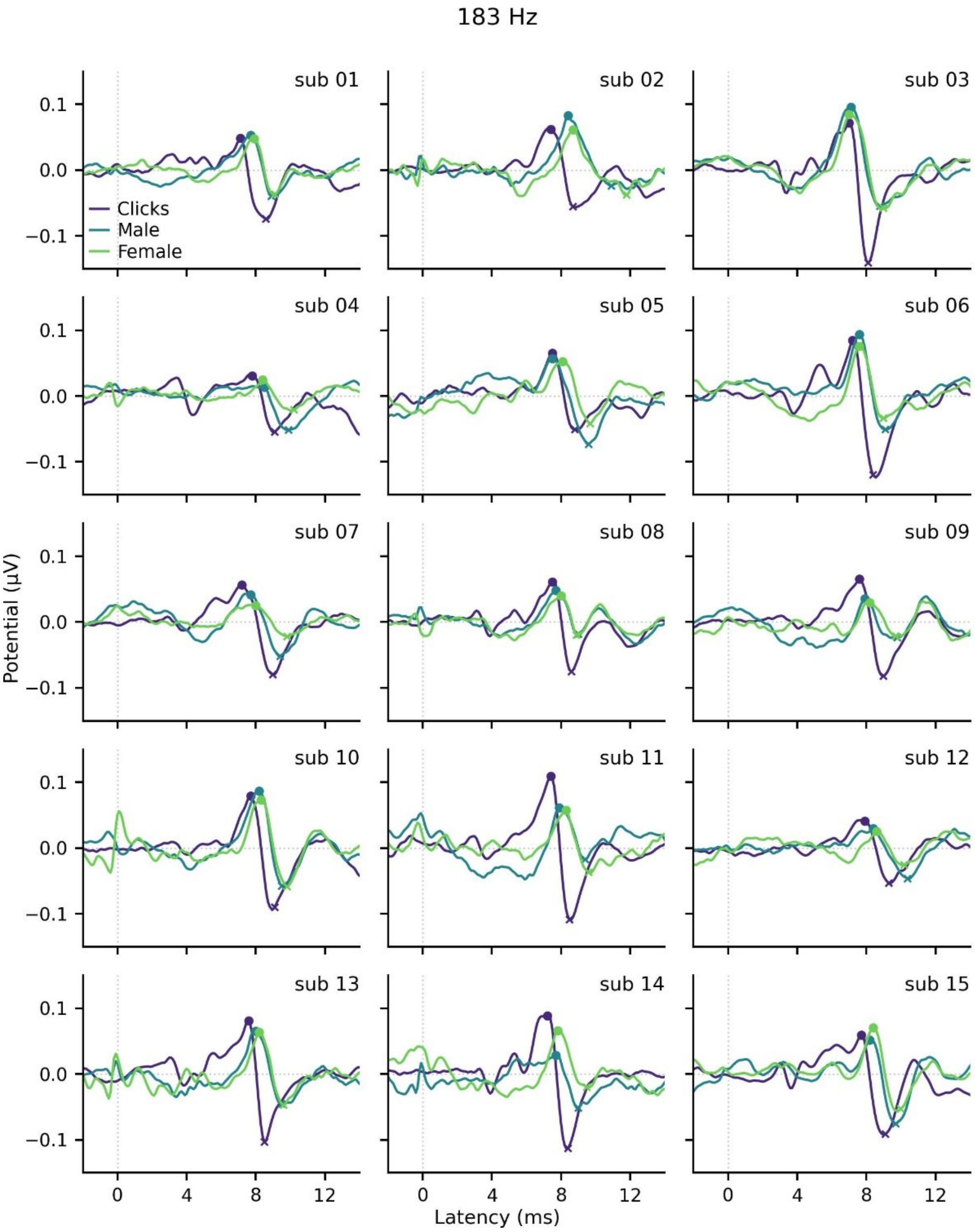


Supplemental Figure 3. Individual participant ABRs are shown for clicks, male- and female-narrated speech with a mean rate or f0 of 183 Hz. Wave V peak latency (circle) and trough (x) are marked, which were used to calculate amplitude.

Supplemental Table 1. Linear mixed effects model for wave V peak amplitude.

Formula[a]: amplitude (nV) ~ stimulus + Hz + stimulus:Hz + (1 | participant)

| | Estimate ± SE | Df | t | p |
|---|---|---|---|---|
| Intercept (default: male) | 132.66 ± 9.13 | 17.43 | 14.53 | **<0.001** |
| Female | -16.26 ± 5.10 | 115.00 | -3.19 | **0.002** |
| Clicks | 35.10 ± 5.10 | 115.00 | 6.88 | **<0.001** |
| Hz (default: male) | -0.81 ± 0.15 | 115.00 | -5.54 | **<0.001** |
| Female:Hz | -0.07 ± 0.21 | 115.00 | -0.35 | 0.727 |
| Clicks:Hz | 0.32 ± 0.21 | 115.00 | 1.57 | 0.119 |

[a] Hz was zero-corrected (i.e., 150 subtracted to give -27, 0, 33 Hz)

Supplemental Table 2. Linear mixed effects model for wave V peak latency.

Formula[a]: latency (ms) ~ stimulus + Hz + stimulus:Hz + (1 | participant)

| | Estimate ± SE | Df | t | p |
|---|---|---|---|---|
| Intercept (default: male) | 7.76 ± 0.09 | 15.49 | 91.14 | **<0.001** |
| Female | 0.17 ± 0.03 | 115.00 | 5.30 | **<0.001** |
| Clicks | -0.35 ± 0.03 | 115.00 | -10.78 | **<0.001** |
| Hz (default: male) | 0.0039 ± 0.0009 | 115.00 | 4.17 | **<0.001** |
| Female:Hz | 0.0011 ± 0.0013 | 115.00 | 0.81 | 0.419 |
| Clicks:Hz | -0.0025 ± 0.0013 | 115.00 | -1.87 | 0.064 |

[a] Hz was zero-corrected (i.e., 150 subtracted to give -27, 0, 33 Hz)
